# RNA3D-Decoy: A Large-Scale Benchmark Dataset of AI-Generated RNA Structure Models for Quality Assessment

**DOI:** 10.64898/2026.10.07.757332

**Authors:** Xiao Chen, Hpoo Yadi, Hanyu Liu, Renzhi Cao

**Affiliations:** Computer science department, Hamilton College, Clinton, NY, USA; Computer science department, Pacific Lutheran University, WA, USA

**Keywords:** RNA, Quality assessment, Benchmark

## Abstract

Recent advances in AI-based RNA structure prediction have increased the availability of candidate models, but their accuracy remains variable. Developing reliable quality assessment (QA) methods requires datasets that represent contemporary prediction errors across diverse targets and molecular contexts. Here, we present RNA3D-Decoy, a large-scale benchmark dataset comprising 103,549 decoys for 677 experimentally determined targets, including 416 RNA-only structures, 117 RNA structures with metal ions, and 144 RNA-protein complexes. Decoys were generated using five biomolecular predictors: AlphaFold 3, ESMFold2, Protenix, Chai-1, and Boltz-2. Each decoy is evaluated against its experimental reference using TM-score, lDDT, and RMSD, with global and local quality annotations where applicable. DockQ scores additionally characterize RNA-protein interface quality. Target grouping incorporates both RNA and protein sequence relationships to support dataset partitioning. Analyses reveal distinct quality distributions across dataset levels and target-dependent differences among predictors, with each predictor contributing the highest-TM-score model for a subset of targets. Broad pooled score distributions also coexist with limited within-target score variation, particularly for RNA-protein complexes, emphasizing the importance of target-level characterization. By combining AI-generated decoys, standardized quality labels, and structured metadata, RNA3D-Decoy provides a resource for developing and evaluating methods for global quality estimation, local error prediction, and candidate-model ranking. The dataset and accompanying code are available at: https://github.com/HamiltonChenLab/RNA3D-Decoy.

## Introduction

The three-dimensional structure of RNA governs its molecular recognition, catalytic activity, regulatory function, and interactions with proteins, ligands, and ions. Experimental techniques, including X-ray crystallography, cryogenic electron microscopy, and nuclear magnetic resonance spectroscopy, remain the primary sources of high-resolution RNA structures. However, RNA flexibility, conformational heterogeneity, and dependence on the molecular environment can complicate experimental characterization. Structure determination is also costly and time-consuming, leaving a substantial gap between the number of known RNA sequences and the number of experimentally resolved RNA structures.

Computational approaches have long provided a complementary route to RNA structure determination. Earlier methods relied primarily on comparative modeling, fragment assembly, conformational sampling, and energy-based optimization. More recent deep-learning methods learn structural constraints or atomic coordinates directly from sequence-derived information. For example, trRosettaRNA2(1) predicts inter-nucleotide geometric relationships and uses them to guide three-dimensional structure construction, while DeepFoldRNA integrates multiple-sequence alignments (MSAs) and predicted secondary structures to estimate inter-nucleotide distances, orientations, and torsion angles (2). RhoFold+(3) further incorporates representations learned from large RNA sequence collections to predict RNA tertiary structures. These RNA-focused methods marked a transition from manually specified sampling procedures toward learned structural representations.

Advances in protein structure prediction have also stimulated the development of general-purpose biomolecular modeling systems. Recent methods can model not only isolated RNA chains but also complexes containing proteins, ligands, and ions (3–8). This expansion is important because many RNAs adopt their functional conformations through interactions with other molecular components. Nevertheless, assessments in CASP15 and CASP16 showed that RNA prediction accuracy still varies substantially across targets and predictors (9, 10). Modern predictors also use stochastic sampling and alternative inference settings to generate multiple candidate structures for a single target. Consequently, improved structure generation creates a parallel need for methods that can estimate model accuracy and select reliable structures from heterogeneous candidate pools.

### RNA structure quality assessment

RNA structure quality assessment (QA) methods estimate the accuracy of predicted structures and rank candidate models when the native structure is unavailable. Existing approaches can be broadly divided into three categories. Knowledge-based statistical potentials derive scoring functions from geometric patterns observed in experimentally determined structures (11–14). These methods are generally computationally efficient and do not require large supervised training sets, but their performance depends on the structural representations and statistical potentials selected by the developer.

Consensus-based methods rank models according to their agreement with other candidates in the same decoy pool (15, 16). They can identify a dominant structural cluster without requiring a native reference, but their predictions depend on the composition of the candidate pool. A consensus score may be misleading when most models are inaccurate or when highly correlated predictors generate similar errors.

Deep-learning QA methods instead learn representations of RNA sequence and geometry to predict global or nucleotide-level model quality (17–19). Their ability to learn directly from structural data reduces reliance on manually designed scoring functions, but it also creates a strong dependence on the training set. A useful training resource must contain many independent RNA targets, structurally diverse candidate models, inaccurate patterns representative of the predictors encountered at inference time, and consistently calculated quality labels.

### Existing datasets

Existing RNA QA datasets satisfy only parts of these requirements. Their nominal size is often driven by large numbers of correlated decoys generated for a comparatively small number of experimental targets (18). Consequently, the number of structures may substantially overstate the effective diversity of the training data. Several widely used resources also rely heavily on molecular dynamics, native-structure perturbation, normal-mode sampling, or fragment-assembly protocols. RNA3DCNN, for example, used molecular-dynamics and Rosetta-generated structures, whereas the ARES training set contained FARFAR2 models for only 18 RNAs (17, 18). RNArank expanded target and predictor coverage, but its training targets were restricted to RNAs no longer than 200 nucleotides and its decoy library combined modern predictions with molecular dynamics, backbone perturbation, and fragment-assembly models (19). These resources remain valuable, but their distributions may not fully represent structures produced by current end-to-end biomolecular predictors.

Coverage of molecular context is another limitation. Existing datasets predominantly represent short, isolated RNA chains, whereas ion-dependent RNAs and RNA-protein complexes are less consistently represented. In some aggregated datasets, multichain experimental structures are decomposed into individual chains even though the native conformation may depend on the removed interaction partners. Differences in redundancy control, structural preprocessing, and quality-label definitions further complicate comparisons and dataset integration. Blind community assessments such as RNA-Puzzles and CASP provide realistic and methodologically diverse prediction pools (9, 10, 20), but they include relatively few independent targets and do not provide controlled sampling across predictors. They are therefore well suited to external evaluation but are less suitable as large, standardized resources for supervised QA training.

To address these limitations, we introduce RNA3D-Decoy, a dataset containing 103,549 structure models for 677 experimentally determined targets. Decoys were generated under controlled settings using five end-to-end biomolecular predictors: AlphaFold 3, ESMFold2, Protenix, Chai-1, and Boltz-2. The targets are organized into three levels comprising 416 RNA-only structures, 117 RNA structures containing metal ions, and 144 RNA-protein complexes. Each decoy is evaluated against its experimental structure using TM-score, lDDT, and RMSD, with both global and local quality annotations provided where applicable.

RNA3D-Decoy combines broad target coverage, decoys generated by current AI-based predictors, controlled cross-predictor sampling, and standardized quality labels. The dataset is designed to support the training and evaluation of RNA QA methods for global quality estimation, local error prediction, and candidate-model ranking. Figure 1 summarizes the dataset-construction workflow, and Table 1 compares RNA3D-Decoy with existing RNA model-quality resources.

**Table 1.** RNA 3D structure datasets used for model-quality assessment.

| Dataset/resource | Native RNA targets | Decoy source | Number of decoys |
| --- | --- | --- | --- |
| **RNA3DCNN datasets(17)** | Training: 414 (332 train/82 validation); Test I (RASP): 85; Test II (KB): 20; Test III (RNA-Puzzles Rounds I-III): 18. | Training MD: GROMACS high-temperature simulated annealing; training MC: Rosetta fragment assembly with all-atom refinement. Test I: MODELLER. Test II: restrained MD/REMD and normal-mode perturbation. Test III: blind RNA-Puzzles submissions. | Training MD: 124,200; training Rosetta-MC: 124,200; Test I: 42,500; Test II: 23,950; Test III: 12–70 per target. |
| **ARES datasets (18)** | Train/validation: 18 (14/4); Benchmark 1: 21; Benchmark 2: 16; blind RNA-Puzzles: 4. | FARFAR2. Approximately 1% of Benchmark 1 models used native-coordinate restraints; Benchmark 2 used fully de novo modeling. | Train/validation: 18,000; Benchmark 1: ≈31,500; Benchmark 2: 80,000 corrected decoys. |
| **RNArank training set (19)** | 1,635 experimental RNAs; structures resolved before January 2022; length ≤200 nt after filtering. | Deep-learning prediction, SimRNA, random backbone-angle perturbation, GROMACS/CHARMM36 MD, and Rosetta fragment assembly. | ≈200,000. |
| **RNA-Puzzles(20)** | Rounds I-IV: 22; 2020 toolkit subset: 20; Round V: 23 RNA targets. | Blind community submissions | 2020 toolkit: 972; Rounds I-IV expanded snapshot: 1,028. |
| **CASP15 (9)** | 12 RNA-containing targets: 10 RNA-only and 2 RNA-protein complexes; 8 natural and 4 synthetic. | Blind submissions from 42 predictors. | 1,660 submitted predictions. |
| **CASP16(10)** | 42 nucleic-acid-containing systems, including 35 RNA monomers and 1 DNA monomer; multimer, hybrid, and ligand categories overlap. | Blind submissions from 65 predictors. | ≈6,500 submitted predictions. |
| **RNA3D-Decoy** | Level I: 416; Level II: 117; Level III: 144; Total: 677. | Five end-to-end biomolecular predictors: AF3, ESMFold2, Protenix, Chai-1, and Boltz-2. | Level I: 63,648; Level II: 17,901; Level III: 22,000; Total: 103,549. |

*Abbreviations:* RASP, Ribonucleic Acids Statistical Potential; KB, knowledge-based; MD, molecular dynamics; REMD, replica-exchange molecular dynamics; MC, Monte Carlo. RNA-Puzzles is the benchmark name; counts in the target column refer to RNA targets.

**Fig 1.**
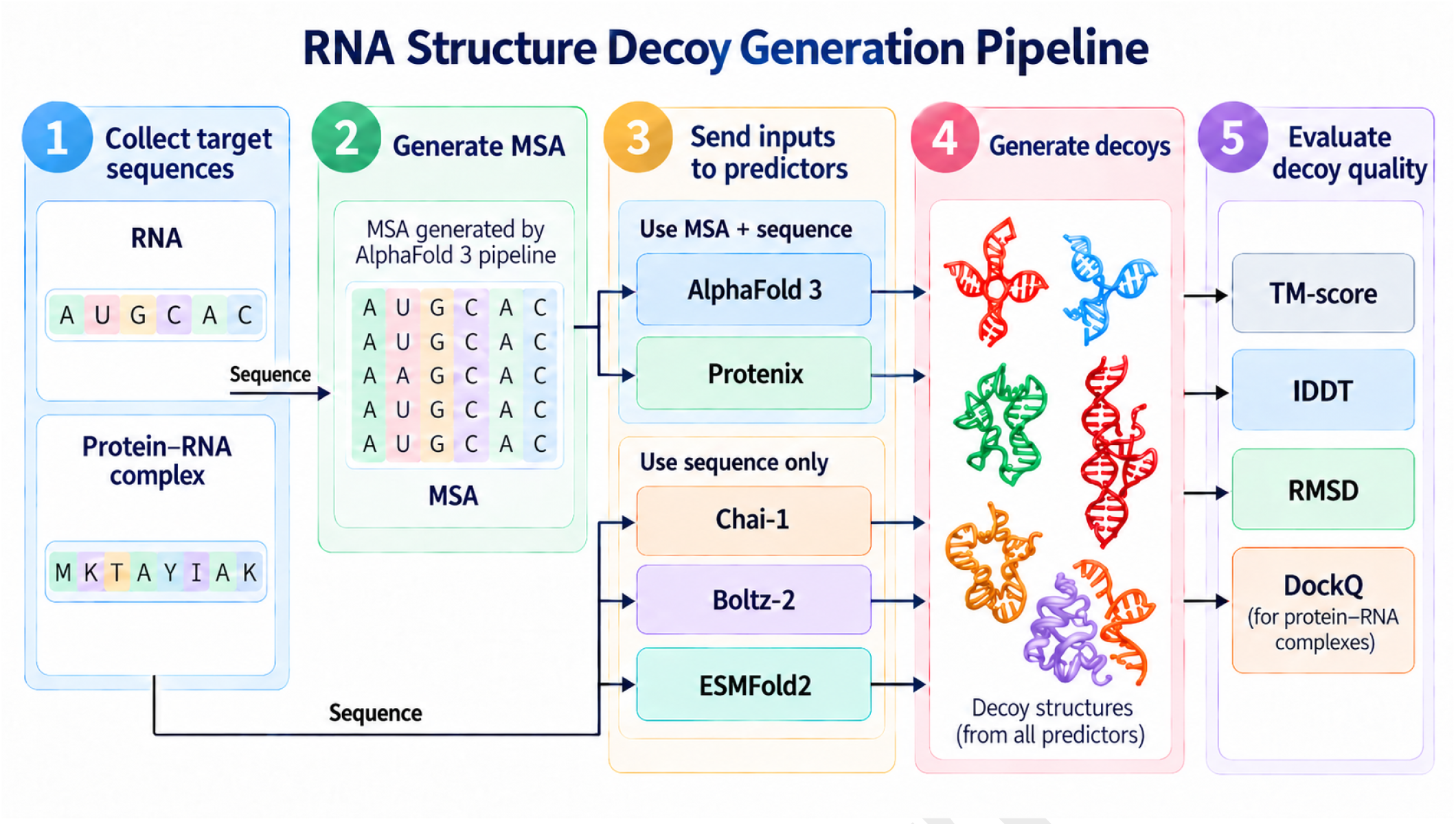
RNA3D-Decoys dataset generation pipeline.

## Materials and Methods

### Dataset design and scope

RNA3D-Decoy was constructed as a standardized collection of experimentally determined RNA structures and computationally generated structural decoys. The dataset was organized into three levels of increasing molecular complexity. Level I contains single-chain RNA structures without explicitly retained interaction partners. Level II contains single-chain RNA structures associated with metal ions. Level III contains RNA-protein complexes comprising exactly one RNA chain and one protein chain. An RNA-protein pair was considered interacting when at least one interatomic distance between the two chains was no greater than 6 Å.

For each target, candidate structures were generated using five end-to-end biomolecular structure predictors: AlphaFold3, ESMFold2, Protenix, Chai-1, and Boltz-2. The experimentally determined structure was used as the native reference. Each generated structure was evaluated using both global and local structural-quality measures. This design enables the dataset to support model-level ranking, residue-level error estimation, and the development of RNA structure quality-assessment methods.

### Collection of experimentally determined RNA targets

Experimentally determined structures were collected from the Protein Data Bank (PDB) using entries available on or before January 1, 2026. Structures determined by X-ray crystallography or cryogenic electron microscopy were required to have a reported resolution of 4.0 Å or better. No resolution threshold was applied to structures determined by nuclear magnetic resonance spectroscopy; for entries containing multiple NMR conformers, only the first deposited model was retained.

Entries containing DNA, additional polymer types, modified RNA residues, or organic ligands were excluded, except for the protein partner required in Level III and the metal ions retained in Level II. Level III targets were further restricted to structures containing exactly one RNA chain and one protein chain satisfying the 6 Å contact criterion.

For Level I, RNA targets with sequences shorter than 10 nucleotides and all of their associated decoys were excluded from the analysis. This minimum-length criterion was applied only to Level I. The retained Level I targets had sequence lengths of 10–720 nucleotides, with a median of 27.5 nucleotides (Table 2). All dataset counts and analyses reported here refer to the retained target set.

**Table 2.**
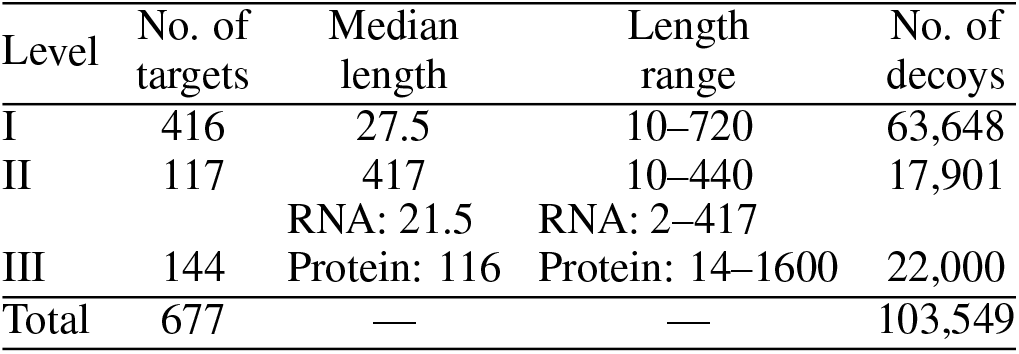
Summary of the retained decoy sets after excluding Level I RNA sequences shorter than 10 nucleotides. RNA lengths are in nucleotides and protein lengths are in amino-acid residues.

| Level | No. of targets | Median length | Length range | No. of decoys |
| --- | --- | --- | --- | --- |
| I | 416 | 27.5 | 10–720 | 63,648 |
| II | 117 | 417 | 10–440 | 17,901 |
|  |  | RNA: 21.5 | RNA: 2–417 |  |
| III | 144 | Protein: 116 | Protein: 14–1600 | 22,000 |
| Total | 677 | — | — | 103,549 |

After applying the level-specific selection criteria, the curated collection contained 416 Level I targets, 117 Level II targets, and 144 Level III targets, for a total of 677 experimentally determined structures. Each retained structure was assigned a stable target identifier linking the native coordinates, input sequence, generated decoys, and quality labels.

### Selection of RNA structure predictors

CASP15 (9) was the first CASP experiment to formally assess RNA three-dimensional structure prediction. Although most RNA-only targets had at least one prediction with the correct global topology, expert-guided, non-deep-learning groups outperformed the submitted deep-learning methods. CASP16 (10) expanded the assessment to RNA and DNA monomers, multimers, and complexes with proteins and small molecules. Automated server performance improved, but the leading groups remained human-expert predictors, and prediction accuracy continued to depend strongly on related experimental templates. Thus, CASP15 and CASP16 highlighted both progress and substantial remaining limitations in automated nucleic-acid structure prediction

To generate decoys representative of the current all-atom prediction regime, we selected AlphaFold3(5), ESMFold2(8), Protenix(7), Chai-1(21), and Boltz-2(22). These methods support RNA-containing inputs, generate all-atom structures, and permit repeated stochastic sampling. AlphaFold 3 introduced a unified Pairformer-and diffusion-based framework for predicting complexes containing proteins, nucleic acids, ligands, ions, and modified residues. ESMFold2 combines ESMC representations with an all-atom diffusion architecture and supports optional MSA conditioning. Protenix is an open-source reproduction of AlphaFold 3 with explicit evaluations on nucleic-acid targets. Chai-1 is a multimodal predictor that supports single-sequence or MSA-assisted inference and optional structural restraints. Boltz-2 extends the open-source Boltz-1 co-folding framework by jointly modeling complex structures and protein–ligand affinity. Collectively, these models represent recent general-purpose biomolecular co-folding systems rather than RNA-specific prediction methods.

### Decoy generation

To predict RNA structures for each target, different structural prediction models such as AlphaFold3 (AF3) (4), ESMFold2 (8), Protenix(7), Chai-1(21) and Boltz-2(22) were used. Multiple decoys were generated for each target by adjusting parameters and settings. Table 3 lists the details for each method.

**Table 3.**
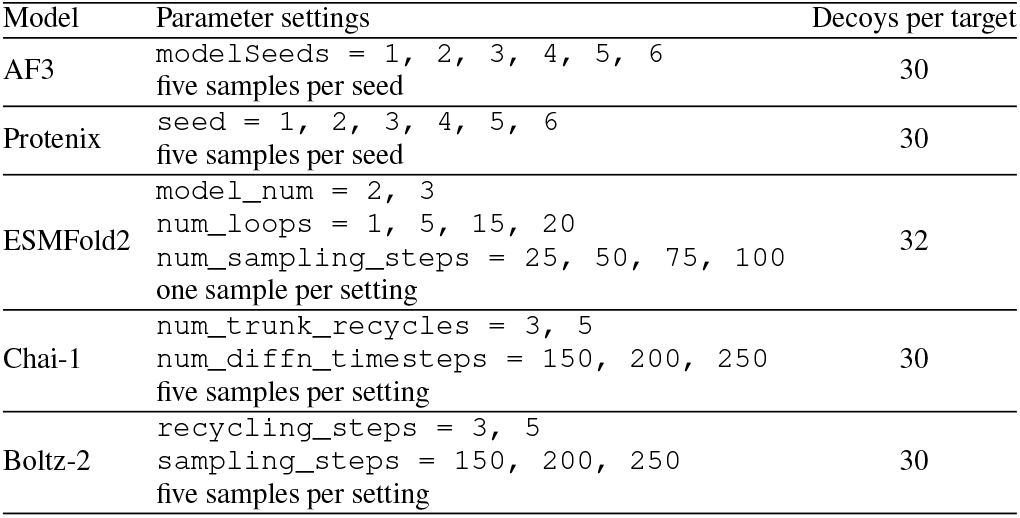
Summary of decoy generation settings for each prediction model. For ESMFold2 model, model_num 2: esmfold2-fast-2026-05, model_num 3: esmfold3-2026-05

MSAs were generated once per target using the Al-phaFold 3 (AF3) data pipeline and reused where supported. Each target FASTA file was converted to AF3 JSON input, and the AF3 data pipeline was run with inference disabled (`-norun_inference`). RNA MSAs were generated with `nhmmer` against the AlphaFold database set: `nt_rna_ 2023_ 02_ 23_clust_seq_id_ 9 0_cov_80_rep_seq. fasta, rfam_14_9_clu st_seq_id_90_cov_80_rep_seq.fasta`, and `rnacentral_active_seq_id_90_cov_80_lin clust.fasta`. For RNA-protein complex targets, protein MSAs were generated with `jackhmmer` against `uniref 90_2022_05.fa, mgy_clusters_2022_05.fa, bfd-first_non_consensus_sequences.fasta`, and `uniprot_all_2021_04.fa`; protein template searches used `pdb_seqres_2022_09_28.fasta` and the PDB mmCIF database. The AF3 data-pipeline outputs were reused for AF3 folding by running inference with the data pipeline disabled (`-norun_data_pipeline`), ensuring that all AF3 seeds used the same precomputed MSAs. For Protenix, the AF3-derived MSA outputs were extracted as A3M files and used as precomputed MSA paths for the corresponding RNA and protein chains. MSAs were not used for ESMFold2, Chai-1, or Boltz-2.

#### A. Global and local quality-label calculation

1. Global and local quality labels were calculated by comparing each generated decoy to its native structure. TM-scores were calculated by USalign(23) with default RNA C3’ atom, while lDDT and RMSD were calculated by OpenStructure v2.11.1 (24). For level 1 and level 2 targets, lDDT and RMSD labels were calculated using RNA C4’ atom selection. For level 3 RNA-protein complexes, lDDT and RMSD labels were calculated using a hybrid atom selection of RNA C4’ atoms and protein CA atoms. Global labels were saved as summary-level scores for each decoy, which represent overall quality of the predicted structure. Local labels were saved as per-atom scores, which represent the quality at individual atom positions.

#### B. Dataset organization and metadata

All generated decoys are stored in the `RNA3D_decoys/` folder. The dataset is organized into three target levels: `level1/` for RNA-only targets, `level2/` for RNA with ions, and `level3/` for RNA-protein complex targets. Each level contains native structures, FASTA files, model-specific decoy folders, and a `labels/` folder that contains the corresponding evaluation metadata. Within each level, decoys are grouped by prediction method: AF3, Protenix, ESMFold2, Chai-1, and Boltz-2. Each `labels/` folder includes TM-score, global and local lDDT, and global and residue-level RMSD files.

**Table 4.**
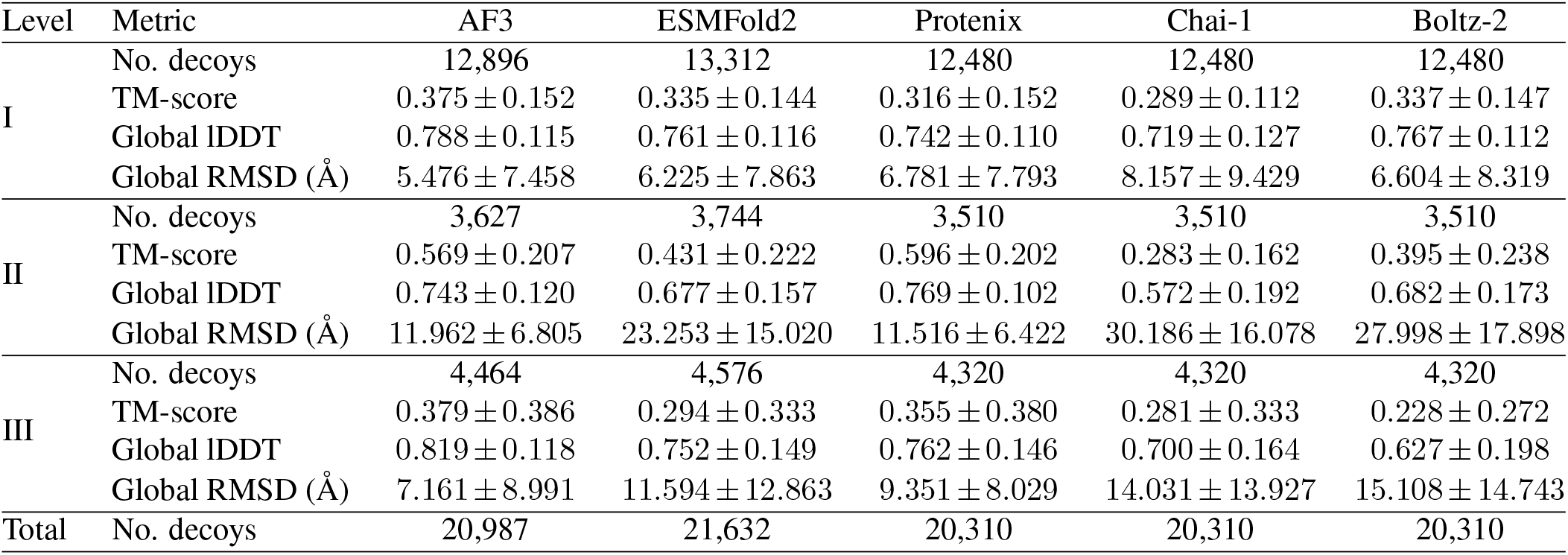
Quality metrics for the 103,549 retained decoys from 677 targets (416, 117, and 144 in Levels I, II, and III, respectively). Scores are reported as mean *±* standard deviation within each method and level. Level I targets with RNA sequences shorter than 10 nucleotides are excluded.

**Table 5.**
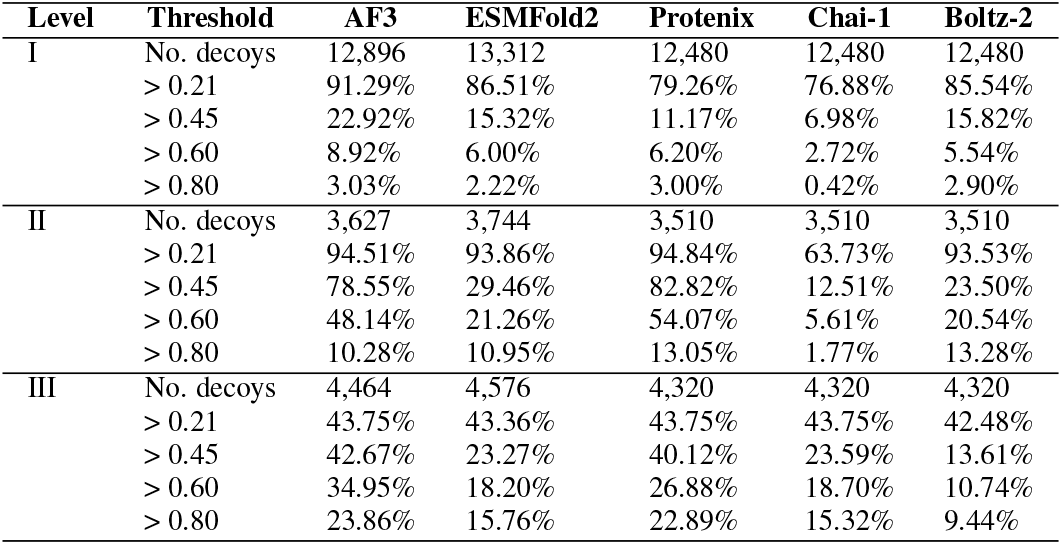
Percentage of decoys with TM-scores strictly greater than 0.21, 0.45, 0.60, and 0.80, calculated separately for each method and level. Panels from left to right show Levels I, II, and III. Levels I and II report RNA scores, and Level III reports RNA-protein complex scores. The thresholds are descriptive cutoffs.

### Sequence clustering

Sequence clustering was performed using the `easy-cluster` workflow in MMseqs2 (25) to group sequence-related targets for dataset partitioning. RNA sequences from Levels 1, 2, and 3 were pooled and clustered jointly, retaining the mapping between each sequence and its source target. RNA clustering was performed in nucleotide mode (`-search-type 3`), with the minimum sequence-identity parameter set to 0.8 (`-min-seq-id 0.8`) and a minimum bidirectional alignment coverage of 0.8 (`-c 0.8-cov-mode 0`). For Level 2, only RNA sequences were included; associated ions did not participate in sequence clustering. Protein sequences from Level 3 complexes were clustered separately using an independent `easy-cluster` run.

The resulting RNA and protein cluster assignments were combined at the target level. Each target was represented as a node in an undirected graph, and two targets were connected if their RNA sequences belonged to the same RNA cluster or, for Level 3 targets, if any of their protein chains belonged to the same protein cluster. The connected components of this graph defined the final target groups, incorporating both cross-level RNA relationships and protein-mediated relationships among Level 3 complexes.

The target lists could be find in the project’s GitHub repository as `train.txt, val.txt`, and `test.txt`.

## Results

In this section, we detailed analyzed all decoys’ TMscore result and Level 3 data’s DockQ result.

### Quality profiles across dataset levels

Structural quality varied across predictors and dataset levels (Table 4). In Level I, AF3 had the highest mean TM-score (0.375 ± 0.152) and global lDDT (0.788 ± 0.115), together with the lowest mean global RMSD (5.476 ± 7.458 Å). Mean TM-scores for the other predictors ranged from 0.289 for Chai-1 to 0.337 for Boltz-2. The ordering of methods was not identical across metrics: ESMFold2 had a lower mean RMSD than Boltz-2 (6.225 versus 6.604 Å), whereas Boltz-2 had slightly higher mean TM-score and global lDDT values.

Protenix had the most favorable mean values for all three metrics in Level II, with a TM-score of 0.596 ± 0.202, global lDDT of 0.769 ± 0.102, and global RMSD of 11.516 ± 6.422 Å. AF3 followed closely, with corresponding values of 0.569 ± 0.207, 0.743 ± 0.120, and 11.962 ± 6.805 Å. ESM-Fold2, Boltz-2, and Chai-1 had lower mean TM-scores of 0.431, 0.395, and 0.283, respectively. Thus, the separation between predictors depended on the target level and the metric used.

For Level III, AF3 again had the highest mean TM-score (0.379 ± 0.386) and global lDDT (0.819 ± 0.118), and the lowest mean RMSD (7.161 ± 8.991 Å). Protenix ranked second on all three mean metrics. These comparisons describe the sampled decoy pools under the generation settings used here. Cross-level differences also reflect distinct target sets and scoring contexts: the TM-score figures report RNA scores for Levels I and II and RNA-protein complex scores for Level III, while the Level III lDDT and RMSD calculations include both RNA and protein atoms.

### TM-score distributions differ across levels

The violin plots and empirical cumulative distribution functions (ECDFs) show that the mean scores summarize markedly different distribution shapes (Figs. 2 and 3). Level I decoys were concentrated at low-to-intermediate TM-scores, with substantial overlap among methods and relatively sparse high-score tails. AF3 showed a shift toward higher scores over much of this distribution. In Level II, AF3 and Protenix had prominent density around a TM-score of 0.6, whereas the main concentrations for the other methods occurred at lower scores. The corresponding ECDFs separated most clearly in the intermediate-score range.

**Fig 2.**
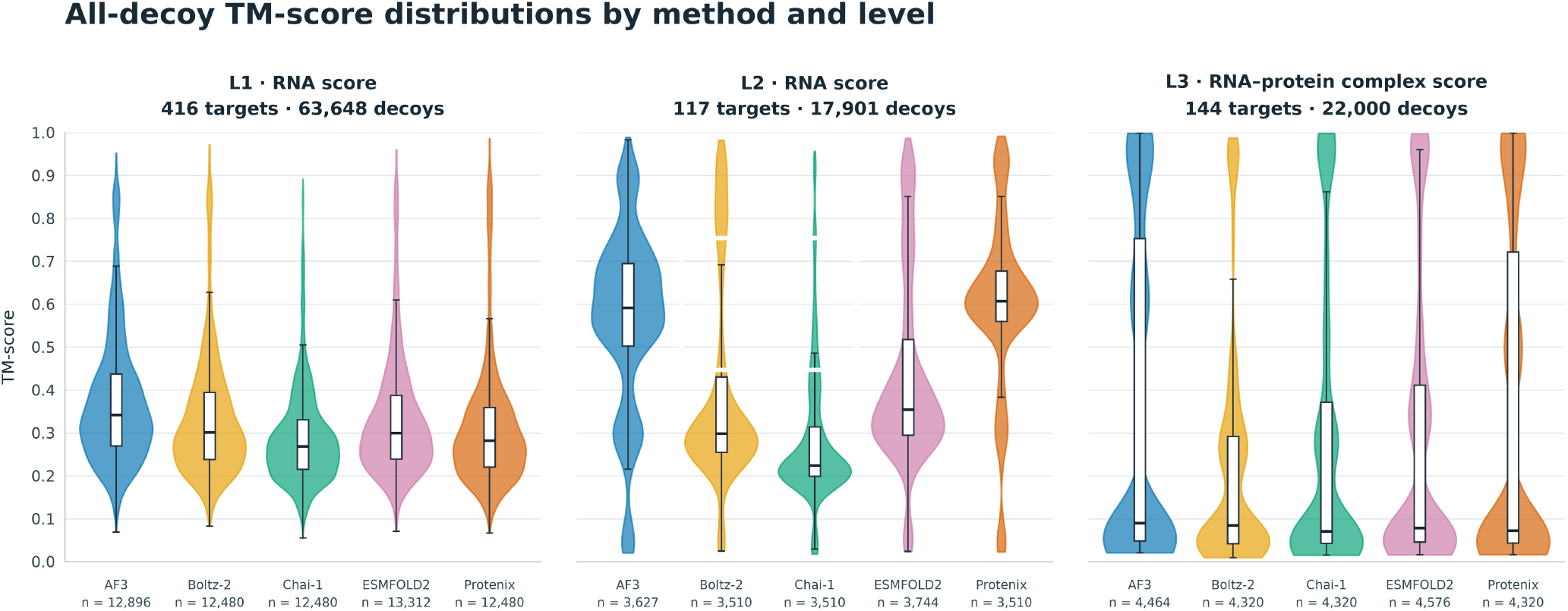
Violin plot for all-decoy TM-score distribution by method and level. From left to right, the panels are level 1 data, level 2 data and level 3 data respectively.

**Fig 3.**
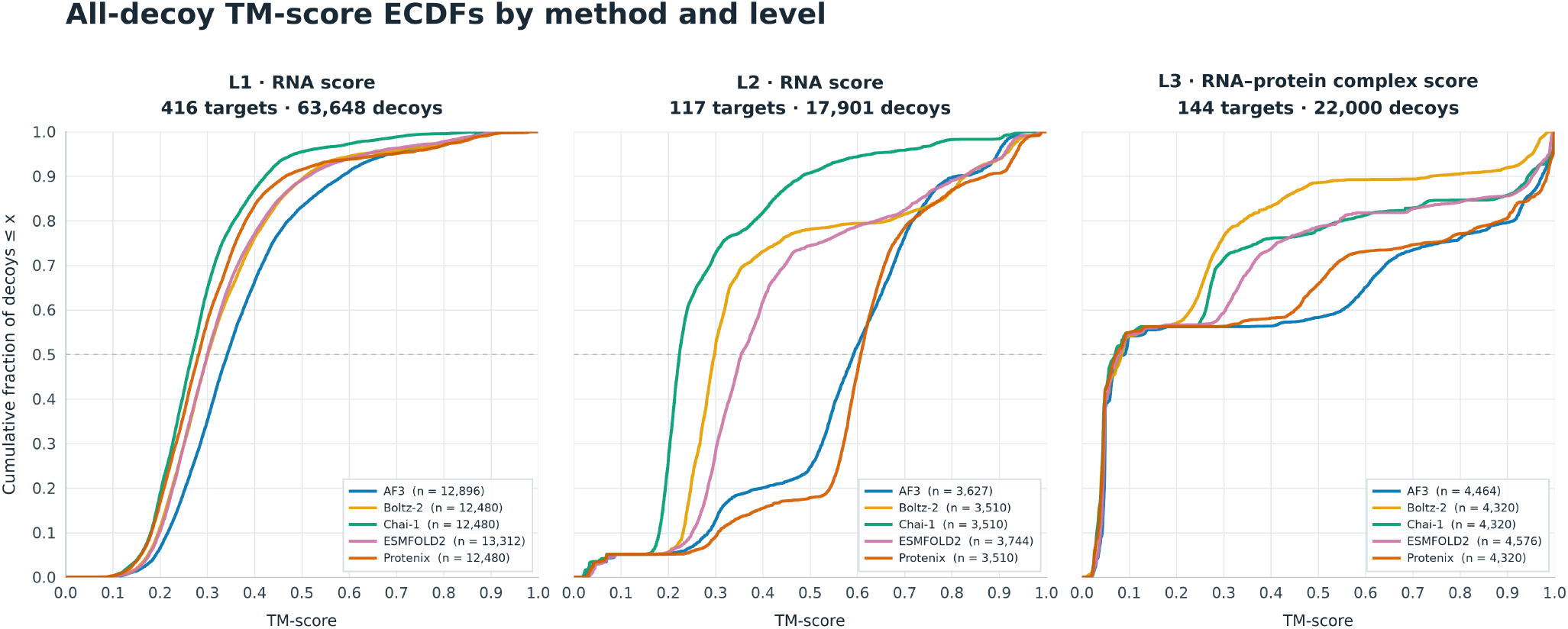
All-decoy TM-score ECDFs by method and level. From left to right, the panels are level 1 data, level 2 data and level 3 data respectively.

Level III showed a pronounced concentration near zero together with a high-score component approaching one. This polarization was present across all five methods, although the relative sizes of the components differed. TM-score standard deviations ranged from 0.272 to 0.386 in Level III, compared with 0.112 to 0.152 in Level I. The broad Level III distributions therefore make the high-score fraction informative alongside the mean.

### TM-score threshold exceedance

The fractions of decoys exceeding fixed TM-score thresholds further distinguished the predictor distributions (Table. **??**). In Level I, AF3 had the largest fraction above 0.45 (22.9%), while the other methods ranged from 7.0% to 15.8%. Only 0.4–3.0% of Level I decoys exceeded 0.8, depending on the predictor. In Level II, 82.8% of Protenix decoys and 78.5% of AF3 decoys exceeded 0.45, compared with 12.5–29.5% for the other methods. At the stricter threshold of 0.8, however, Boltz-2 had the largest reported fraction (13.3%), followed by Protenix (13.0%), despite its lower mean TM-score.

In Level III, AF3 and Protenix had the largest fractions above 0.45 (42.7% and 40.1%) and above 0.8 (23.9% and 22.9%). Across all five methods, only 42.5–43.8% of decoys exceeded 0.21, consistent with the large low-score component visible in the violin plots and ECDFs. These percentages count individual decoys within each method and level. The thresholds serve as descriptive cutoffs, and the reported fractions do not measure the proportion of targets for which at least one high-scoring model was obtained.

### Target-specific rankings and score dispersion

TM-score analysis comprised 103,549 decoys from 677 targets: 416 in Level I, 117 in Level II, and 144 in Level III. Of the 3,385 possible target–method combinations, 3,384 had TM-score summaries. ESMFold2 scores were unavailable for the Level III target 8D97. Direct comparisons among all five predictors therefore used the 676 targets with complete coverage (416, 117, and 143 in Levels I, II, and III, respectively). For each target, we compared both the mean TM-score within each method’s decoy pool and its maximum TM-score. Each target contributed once to each comparison.

AF3 had the highest within-target mean on 410 of 676 targets (60.7%) and the highest maximum on 431 targets (63.8%) (Table 6). ESMFold2 supplied the highest maximum on 102 targets, Boltz-2 on 73, Protenix on 56, and Chai-1 on 14. The target-specific ranking added information to the pooled averages. For example, although Protenix had the highest pooled mean TM-score in Level II, AF3 had the highest within-target mean on 56 of 117 targets (47.9%), compared with 37 targets (31.6%) for Protenix. AF3 also had the highest maximum on 68 Level II targets, compared with 20 for Protenix. Thus, the magnitude of score differences and the number of targets on which a method leads describe different aspects of performance.

**Table 6.** Target-level TM-score comparisons on the 676 targets with results from all five predictors. Counts (percentages) identify the method with the highest within-target mean or maximum. Median IQR summarizes the within-target interquartile range across these targets. No ties occurred at the reported score precision.

| Method | Highest mean | Highest maximum | Median IQR |
| --- | --- | --- | --- |
| AF3 | 410 (60.7%) | 431 (63.8%) | 0.0394 |
| ESMFold2 | 67 (9.9%) | 102 (15.1%) | 0.0441 |
| Protenix | 81 (12.0%) | 56 (8.3%) | 0.0199 |
| Chai-1 | 12 (1.8%) | 14 (2.1%) | 0.0236 |
| Boltz-2 | 106 (15.7%) | 73 (10.8%) | 0.0258 |

Within-target score dispersion also depended on dataset level (Fig. 4). We calculated the interquartile range (IQR) as the 75th minus the 25th percentile within each target–method pool, then took the median across targets. In Level II, median IQRs were 0.1055 for AF3 and 0.0864 for ESMFold2, compared with 0.0279 for Protenix. In Level III, the median IQR was only 0.0026–0.0053 across methods, despite the broad pooled distributions. Moreover, 81 of the 143 fully covered Level III targets (56.6%) had no decoy with a TM-score above 0.21 in any method. These observations indicate that substantial differences between targets coexist with tightly clustered scores within many target–method pools. Score dispersion measures variation in similarity to the reference structure and does not directly measure pairwise structural diversity among decoys.

**Fig 4.**
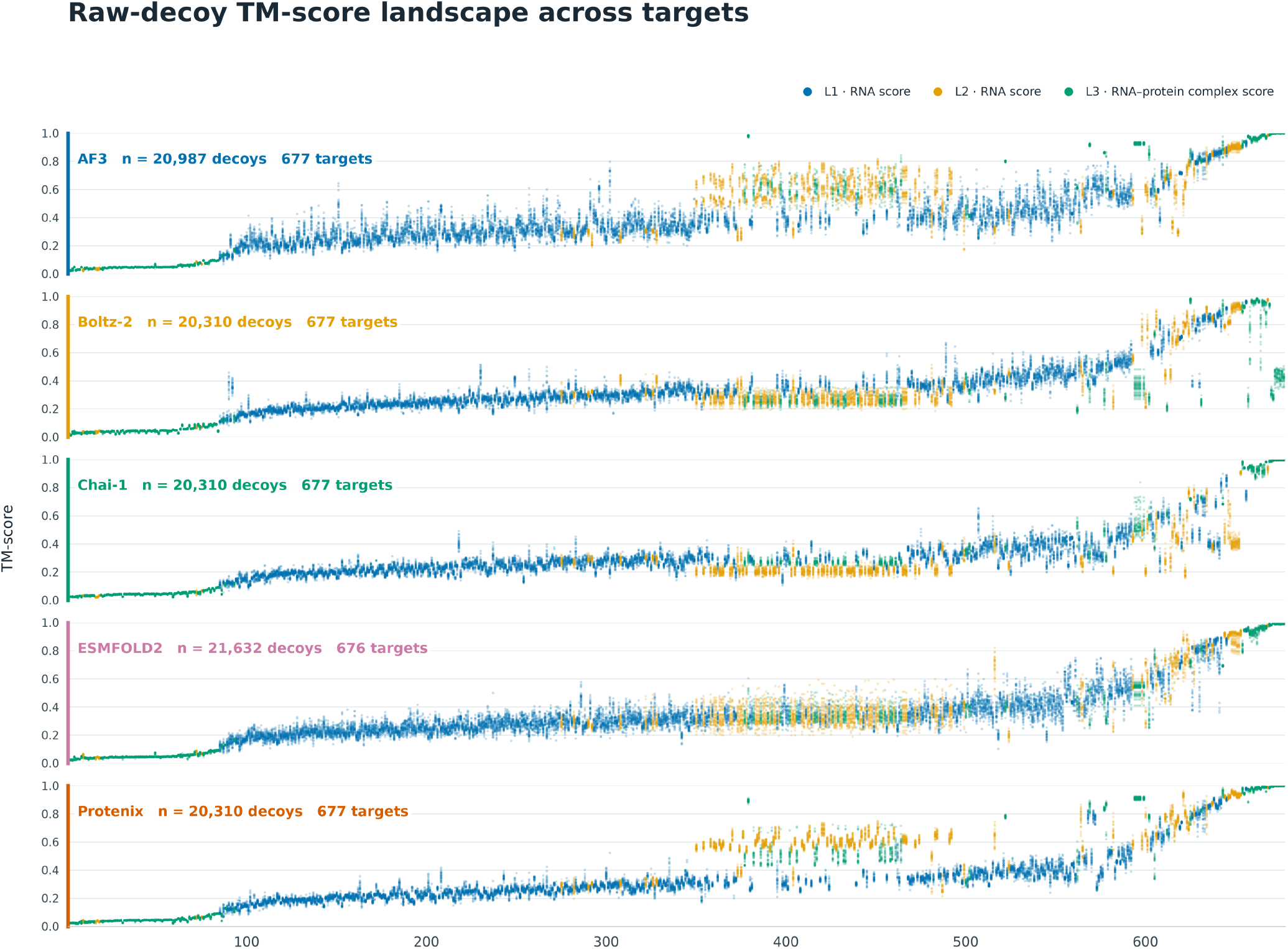
TM-scores of individual decoys across targets. Panels from top to bottom show AF3, Boltz-2, Chai-1, ESMFold2, and Protenix. Colors indicate dataset level. Levels I and II report RNA scores, and Level III reports RNA-protein complex scores.

### Predictor correlations and head-to-head comparisons

The predictor correlations analysis is based on each RNA target’s medain TMscore. According to Fig 5, predictors showed strong correlations in target-level median RMSD for Level I (Spearman ρ = 0.82–0.93), indicating similar rankings of target difficulty. Correlations were more variable in Level II (0.59–0.90) and Level III (0.64–0.90), with the strongest agreement between Chai-1 and Boltz-2, and between Protenix and Chai-1, respectively. Nevertheless, substantial departures from the identity line in the head-to-head plots, particularly in Levels II and III, show that similar target rankings can coexist with considerable differences in absolute prediction error.

**Fig 5.**
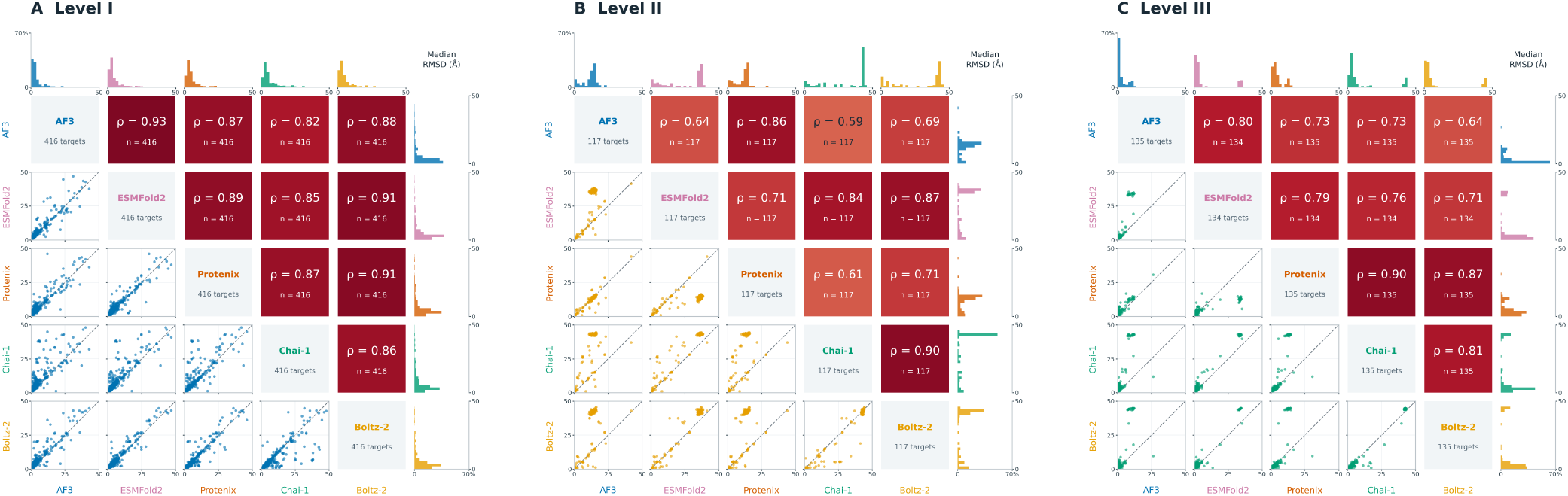
Predictor correlation and head-to-head comparisons plot

### DockQ score threshold exceedance

Table 7 shows across 22,000 Level 3 decoys, AF3 achieved the highest proportions ofmedium-or-higher quality predictions (59.18%) and high-quality predictions (23.10%) under the specified DockQ reference thresholds. ESMFOLD2 ranked second in medium-or-higher coverage (44.17%), although Protenix produced a larger high-quality fraction than ESM-FOLD2 (13.94% versus 10.62%). Overall, 41.67% of decoys were classified as incorrect, indicating substantial remaining limitations in RNA-protein interface prediction within this dataset.

**Table 7.** All-decoy DockQ reference-quality distribution for Level 3. *D* denotes DockQ.

| Method | No. decoys | Incorrect<br>$D < 0.23$ | Acceptable<br>$0.23 \leq D < 0.49$ | Medium<br>$0.49 \leq D < 0.80$ | High<br>$D \geq 0.80$ |
| --- | --- | --- | --- | --- | --- |
| AF3 | 4,464 | 1,168 (26.16%) | 654 (14.65%) | 1,611 (36.09%) | 1,031 (23.10%) |
| ESMFOLD2 | 4,576 | 1,510 (33.00%) | 1,045 (22.84%) | 1,535 (33.54%) | 486 (10.62%) |
| Protenix | 4,320 | 2,193 (50.76%) | 883 (20.44%) | 642 (14.86%) | 602 (13.94%) |
| Chai-1 | 4,320 | 2,365 (54.75%) | 1,139 (26.37%) | 500 (11.57%) | 316 (7.31%) |
| Boltz-2 | 4,320 | 1,931 (44.70%) | 1,177 (27.25%) | 1,037 (24.00%) | 175 (4.05%) |
| Total | 22,000 | 9,167 (41.67%) | 4,898 (22.26%) | 5,325 (24.20%) | 2,610 (11.86%) |

## Conclusions

We present RNA3D-Decoy, a large-scale dataset comprising 103,549 AI-generated decoys for 677 experimentally determined targets. The dataset covers RNA-only structures, RNA structures with metal ions, and RNA-protein complexes, with decoys generated by AlphaFold 3, ESMFold2, Protenix, Chai-1, and Boltz-2. Each decoy is evaluated against its experimental reference using TM-score, lDDT, and RMSD, with global and local annotations where applicable and additional DockQ scores for RNA-protein complexes. This combination extends RNA quality-assessment resources beyond isolated RNA chains and captures prediction errors produced by multiple contemporary biomolecular predictors.

Our analyses revealed substantial variation in model quality across predictors, targets, and dataset levels. AlphaFold 3 most frequently produced the highest-TM-score decoy in target-level comparisons, but each predictor contributed the highest-scoring model for a subset of targets. Broad pooled score distributions also coexisted with limited within-target score variation, particularly in Level III, highlighting the importance of target-level analyses alongside aggregate statistics. DockQ results further showed considerable variability in RNA-protein interface quality, supporting the inclusion of interface-specific annotations alongside overall structural similarity measures. These observations characterize the sampled decoy pools under the generation settings used in this study, rather than establish a definitive ranking of the predictors.

By combining AI-generated decoys with standardized quality labels, structured metadata, and target grouping based on both RNA and protein sequence relationships, RNA3D-Decoy provides a foundation for developing and evaluating RNA QA methods. The dataset is intended to support global quality estimation, local error prediction, and candidate-model ranking across different molecular contexts, while enabling systematic investigation of the strengths and limitations of quality assessment for modern RNA structure predictions.

## ACKNOWLEDGEMENTS

We appreciate RCSB’s public dataset.

---

**Supplementary Note 1: Detailed data statistics**

The supplementary.xlsx Sheet1 describes each target’s sequence length, resolution, method of experimental structure, RNA classification type and Organism information. Sheet2 describes level-2’s target’s ion information. These information has been used when generating level-2 decoys.Sheet3 and Sheet4 describe the aggregated information for global lDDT and global RMSD respectively.

